# HepaCAM Suppresses Glioblastoma Stem Cell Invasion in the Brain

**DOI:** 10.1101/2022.08.24.504654

**Authors:** Arpan De, John M. Lattier, John E. Morales, Jack R. Kelly, Xiaofeng Zheng, Zhihua Chen, Sumod Sebastian, Jason T. Huse, Frederick F. Lang, Joseph H. McCarty

## Abstract

Glioblastoma (GBM) is a malignant brain cancer that contains sub-populations of highly invasive tumor cells that drive progression and recurrence after surgery and radiochemotherapy. The exact mechanisms that enable GBM cells to disperse from the main tumor mass and navigate throughout the brain microenvironment remain largely unknown. As a result, there is a lack of effective strategies to block cancer cell invasive growth in primary and recurrent GBM. Here we report that hepatocyte cell adhesion molecule (hepaCAM), which is normally expressed in perivascular astrocytes, plays central roles in controlling the invasive growth features of GBM cells. Genetically targeting HEPACAM induces a transition from GBM cell proliferation/self-renewal to invasion. Increased invasion is due, in part, to an activation of focal adhesion signaling pathways and enhanced GBM cell adhesion to the extracellular matrix (ECM) in the brain microenvironment. Transcriptional profiling of GBM cells reveals various HEPACAM-regulated genes with links to polarity and invasion. Collectively, these data show that hepaCAM balances ECM adhesion and signaling pathways to control cancer cell proliferation versus invasion in the brain parenchyma. Targeting select components of the hepaCAM pathway may be an effective way to block tumor progression and recurrence in patients with GBM.

## Introduction

HepaCAM, also known as glial cell adhesion molecule (GlialCAM), is a 50-kDa singlepass type I transmembrane glycoprotein with two extracellular immunoglobulin (IgG)-like domains and a ~125 amino acid cytoplasmic tail, that is predominantly expressed in the brain and liver (1). HepaCAM in the liver suppresses the growth of hepatocytes and is down-regulated in hepatocellular carcinoma, suggesting tumor suppressor-like functions (2). In the brain, hepaCAM is normally expressed in astrocytes where it regulates ion homeostasis (3), BBB physiology (4), and synaptic excitation (5). Putative functions for hepaCAM in the brain were discovered during genomic sequencing of patients with the neurodevelopmental brain disorder megalencephalic leukoencephalopathy with subcortical cysts (MLC) (6). MLC patients contain recessive mutations in the HEPACAM gene or in the MLC1 gene, which encodes a 38-kDa protein with 8 transmembrane domains (7). The extracellular IgG-like domains of hepaCAM mediate interactions with Mlc1 as well as other proteins such as the chloride channel ClC-2 (8) and components of the dystrophin-glycoprotein (DGC) complex including aquaporin-4 (9).

We have reported previously that Mlc1 is essential for promoting tumor cell growth and invasion in the brain microenvironment (10). Mlc1 protein is overexpressed in human GBM, a malignant brain cancer that contains sub-populations of proliferative and invasive cells that drive tumor growth, progression and recurrence after therapy (11). MLC1 is a molecular marker for the classical GBM sub-type which is defined in part by EGFR overexpression and wild type TP53 status (12). Silencing MLC1 expression in human GBM stem cells (GSCs) leads to defective self-renewal in vitro and impaired invasion in pre-clinical mouse models. Reduced Mlc1 protein expression in GSCs is associated with the hyperactivation of receptor tyrosine kinase (RTK) signaling pathways, particularly those involving Axl.

Roles for HEPACAM in brain tumors and its possible links to regulation of MLC1 functions, however, have remained largely unexplored. Here, we have analyzed functions for hepaCAM adhesion and signaling pathways in human GBM using a blend of human patient samples and pre-clinical mouse models. We report that hepaCAM coordinately promotes GBM cell proliferation and inhibits invasion in the brain. Genetic inhibition of HEPACAM leads to reduced GBM cell proliferation and self-renewal but increased invasion in vitro and in vivo. HepaCAM control of GBM cell invasion is linked to activation of focal adhesion signaling dynamics as well as increased adhesion to the ECM via integrins. We have also identified various HEPACAM-regulated genes with functions in GSC adhesion and invasion. In summary, these data reveal that hepaCAM modulates GBM cell growth and invasion in the brain microenvironment and suggest that adhesion and signaling effectors regulated by hepaCAM may be attractive therapeutic targets to benefit patients with GBM.

## Results

### HEPACAM is differentially expressed in GBM versus non-cancerous brain tissue

To determine potential roles for HEPACAM in GBM initiation and/or progression, we first queried The Cancer Genome Atlas (TCGA) database for relative levels of HEPACAM mRNA in GBM tissues and non-cancerous brain. As shown in Fig. 1A, significant increases in expression of HEPACAM mRNA were found in GBM samples. In support, immunoblotting detergent-soluble protein lysates prepared from freshly resected GBM samples (n=8) revealed robust hepaCAM protein expression in 7 of 8 samples (Fig. 1B). We also performed immunohistochemical staining to analyze spatial patterns of hepaCAM protein expression in fixed normal human brain and GBM tissue samples. Analysis of regions in the normal human cortex revealed enriched hepaCAM protein levels in astrocytes surrounding cerebral blood vessels (Fig. 1C). Anti-hepaCAM immunostaining of different primary GBM samples taken from non-necrotic tumor regions revealed robust expression in cancer cells throughout the tumor core as well as enrichment in perivascular tumor cells (Fig. 1D, E and Supplemental Fig. 1). The specificity of the anti-hepaCAM antibody was validated by staining fixed human tumor sections with species-matched IgG control and by immunoblotting lysates before and after PNGase treatment (Supplemental Fig. 1). Several genes showing co-expression with HEPACAM in non-cancerous human brain samples and in GBM tissue samples were identified using Correlation AnalyzeR (13) and the ARCHS4 genomic platform (14), revealing putative links between HEPACAM and genes with established roles in brain tumor biology (Supplemental Fig. 2). Querying the Brain RNAseq database (brainrnaseq.org), which contains quantitative gene expression data for different neural and vascular cell types isolated from the neonatal mouse brain, revealed that Hepacam mRNA is highly enriched in astrocytes (Supplemental Fig. 3). Similarly, in an adult mouse brain database (betsholtzlab.org/VascularSingleCells/database.html), Hepacam mRNA expression was detected in astrocytes as well as in oligodendrocytes (Supplemental Fig. 3). Furthermore, HEPACAM mRNA is highly expressed in astrocytes and oligodendroglia of the human brain (Supplemental Fig. 3) as revealed by querying the human BBB database (twc-stanford.shinyapps.io/human_bbb) (15). These results match with our immunohistochemical data from non-cancerous human brain tissue sections that showed enriched hepaCAM protein expression in astrocytes adjacent to blood vessels (Fig. 1C). There were varying levels of HEPACAM mRNA in different regions of non-cancerous mouse and human brain samples with highest expression in the cerebral cortex (Supplemental Fig. 4. Subsequent analysis of the TCGA database for cancers across multiple organs, confirmed elevated levels of HEPACAM mRNA mainly in gliomas (Supplemental Fig. 5) which matches the immunohistochemistry results for hepaCAM protein in fixed GBM samples (Fig. 1C-E and Supplemental Fig. 1).

**Figure 1.**
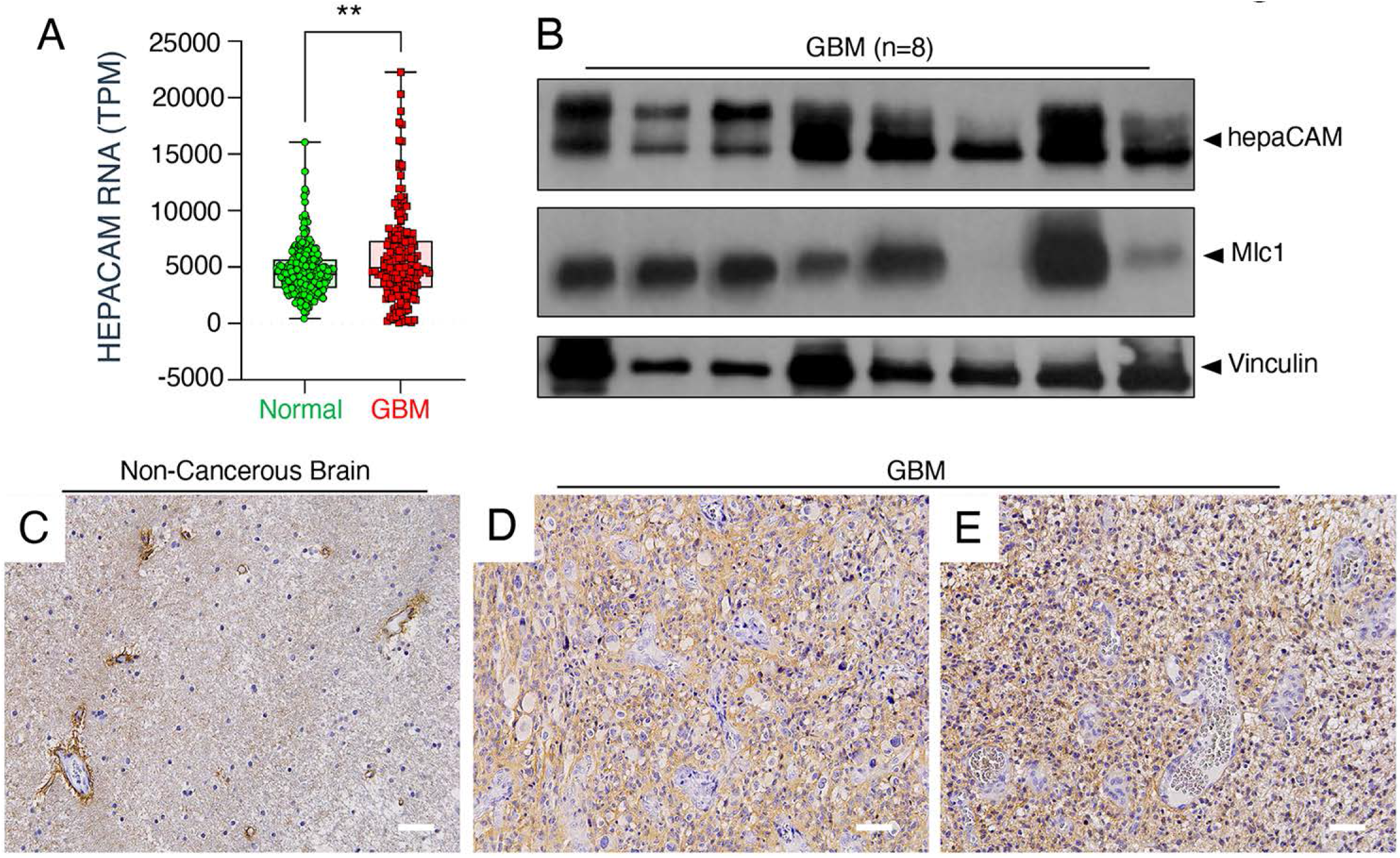
Analysis of HEPACAM expression in the non-cancerous brain versus GBM. **(A);** Querying the GTex and TCGA GBM databases reveals that HEPACAM mRNA levels are significantly higher in GBM (n=173) versus non-cancerous brain (n=207), **p<0.01. **(B);** Anti-hepaCAM immunoblot analysis of a panel of 8 different detergent-soluble lysates from freshly resected human GBM tissue. Note that all GBM samples express detectable levels of hepaCAM protein. **(C-E);** Anti-hepaCAM immunohistochemical staining of fixed non-cancerous human brain (C) and two different resected human GBM tissue samples (D, E). Note the enrichment of hepaCAM protein surrounding blood vessels in the normal brain, whereas there is broader expression of hepaCAM in tumor cells within the GBMs. Scale bars, 50 μm.

In comparison to primary GBM, tumors that recur after surgery and chemotherapy tend to be more infiltrative and invasive (16). Therefore, we analyzed HEPACAM expression levels in samples of the Chinese Glioma Genome Atlas (CGGA), which contains transcriptomic data from a large number of primary and recurrent glioma samples (17). As shown in Supplemental Fig. 6, in comparison to primary glioma surgical samples, there is a significant reduction in HEPACAM mRNA levels in recurrent tumors. The data for diminished HEPACAM levels in recurrent GBM samples match with reduced HEPACAM mRNA levels in low grade gliomas versus non-cancerous human brain samples (Supplemental Fig. 7). Low grade gliomas are diffusely infiltrative and invasive but not highly proliferative like grade III astrocytomas or GBM (18), supporting the hypothesis that down-regulation of HEPACAM gene expression is linked to increased tumor cell invasion.

### Genetic inhibition of HEPACAM leads to reduced GSC growth and enhanced invasion in vitro

To further characterize functions for HEPACAM in GBM, we analyzed expression in primary GBM stem cells (GSCs), which have cancer-initiating capacities and drive tumor progression and recurrence after therapy (19). Robust levels of hepaCAM protein were detected in 7 of 8 primary human GSC spheroid samples analyzed (Fig. 2A). We selected GSC6-27 spheroids for more detailed mechanistic analyses of HEPACAM functions, since these cells express robust levels of hepaCAM protein (Fig. 2A) and generate proliferative and infiltrative tumors after implantation in xenograft mice (20). GSCs were stably infected with pGIPZ lentiviruses expressing GFP and shRNAs targeting HEPACAM (n = 3). pGIPZ lentivirus expressing GFP and non-targeting (NT) shRNAs was used as a control. All three shRNAs targeting HEPACAM resulted in reduced protein expression (30-70%) as determined by immunoblotting detergent-soluble cell lysates (Fig. 2B) or immunofluorescence labeling of GSCs (Fig. 2C) with anti-hepaCAM antibodies. GSC6-27 cells stably expressing shRNA #3, which generated a 70% reduction of hepaCAM expression, were selected for subsequent functional studies. Sphere formation assays were performed to test for hepaCAM-dependent differences in GBM cell proliferation and/or self-renewal. Analysis of spheroid formation over a 7-day period revealed a major reduction in growth capacities of GBM cells expressing HEPACAM shRNAs (Fig. 2D). In addition, spheroid sizes were significantly reduced in HEPACAM shRNA samples versus NT shRNA cells (Fig. 2E, F). After 7 days in culture, cross-sectional areas of nearly 40% of spheroids expressing control NT shRNAs were 2 × 10^4^ μm^2^ or greater compared to only 15% of spheroids expressing HEPACAM shRNAs. Silencing HEPACAM in two additional primary GBM spheroid cultures (GSC231 and GSC2-14) using pGIPZ lentiviral-expressed shRNAs resulted in similar defects in sphere formation (Supplemental Fig. 8). Next, we investigated functions for hepaCAM in GSC invasion in vitro (21). Spheroids were dissociated and cells were exposed to a serum gradient to induce directional invasion through an ECM-coated transwell. In comparison to GSC6-27 cells expressing pGIPZ-infected control (NT), cells expressing HEPACAM shRNAs showed a two-fold increase in invasive capacities (Fig. 2G). GSC231 and GSC2-14 cells also showed similar hepaCAM-dependent increases in invasion through three-dimensional ECM (Supplemental Fig. 8).

**Figure 2.**
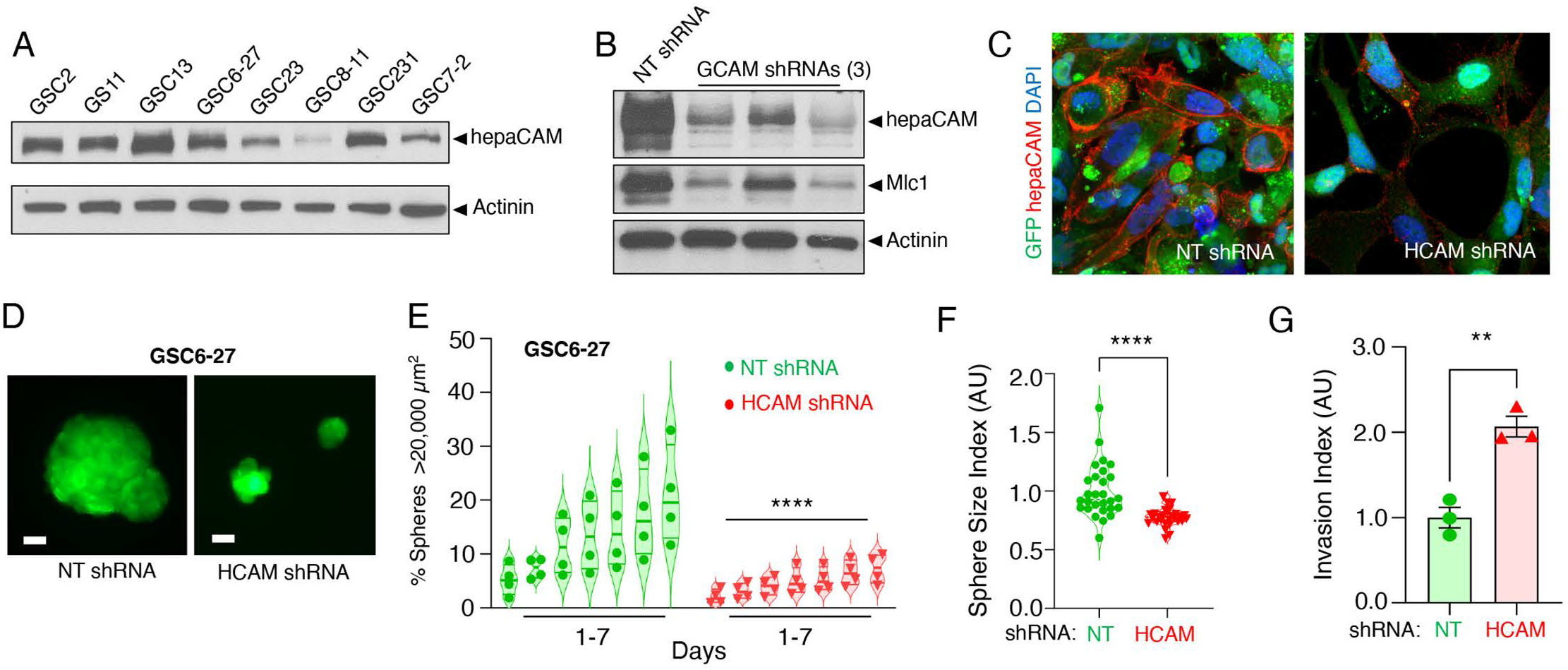
HEPACAM promotes GSC proliferation and self-renewal. **(A);** Detergent-soluble lysates from eight different primary human GSC spheroid cultures were immunoblotted with anti-hepaCAM antibodies. Note the robust expression of hepaCAM protein in 7 of 8 GSC samples. **(B);** pGIPZ lentivirus-expressed shRNAs were tested for HEPACAM (n=3) gene silencing in infected GSC6-27 cells. The different HEPACAM shRNAs silenced gene expression to varying levels (30-70%) as determined by anti-hepaCAM immunoblotting. Note that expression of HEPACAM shRNAs result in slightly diminished levels of Mlc1 protein. As controls for these experiments we used GSC6-27 cells infected with pGIPZ expressing non-targeting (NT) shRNAs. **(C);** GFP+GSC6-27 cells expressing NT control shRNAs (left) or HEPACAM shRNAs (right) were labeled with anti-hepaCAM antibodies. Note the decrease in hepaCAM protein expression (red) in cells following shRNA-mediated gene silencing. **(D);**GSC6-27 cells infected with pGIPZ lentivirus expressing control NT shRNAs formed spheroids that were much larger by seven days in culture than spheroids formed from cells expressing HEPACAM shRNAs (shRNA #3 in panel B). **(E);** The percentages of newly formed spheres (>20,000 μm^2^) were recorded daily for 7 consecutive days. GSC6-27 cells expressing HEPACAM shRNAs showed obviously reduced spheroid formation with significantly smaller cross-sectional areas (ANOVA test and Tukey post-hoc analysis for comparison, n=4, ****p<0.0001) **(F);** Single cell suspensions were allowed to form spheroids over 7 days. The diameter of newly formed spheroids was daily recorded, revealing that spheroids with reduced hepaCAM were significantly smaller in comparison to control GSC6-27 cells that express hepaCAM(Students t-test for comparison, n=4, ****p<0.0001) **(G);** GSC6-27 cells expressing HEPACAM shRNAs showed enhanced invasion through basement membrane-coated transwells as compared to NT shRNA control cells (Student’s t-test for comparison, n=3, **p<0.01).

### Genetic inhibition of HEPACAM leads to enhanced GBM cell invasion in vivo

We next analyzed the functions for hepaCAM in tumor growth and invasion in vivo. GSC6-27 cells infected with pGIPZ virus expressing GFP and NT shRNAs (n = 5) or HEPACAM shRNAs (n = 9) were intracranially injected into the striatum of NCR nu/nu mice. Animals were monitored for tumor-induced neurological deficits over a 16-week time period. When the first animal showed deficits all mice were sacrificed. Anti-GFP antibodies were used to immunolabel fixed brain sections to monitor tumor growth and invasion. Analysis of GFP expression patterns revealed increased numbers of cells expressing HEPACAM shRNAs in the corpus callosum and contralateral (non-injected) hemisphere, as compared to cells expressing NT shRNAs (Fig. 3A-H). On average, 254 GFP^+^ cells were found in the corpus callosum and 182 GFP^+^ cells were found in the contralateral hemispheres of mice injected with GSCs expressing HEPACAM shRNAs, as compared to 101 and 48 GFP^+^ cells in the same brain regions, respectively, of mice injected with cells expressing NT shRNAs. Quantitation of GFP intensity in the corpus callosum revealed hepaCAM-dependent increases in cell invasion as well as differences in GBM cell shape (Fig. 3I, J). GBM cells expressing HEPACAM shRNAs showed robust perivascular invasion near the corpus callosum and more infiltrative growth in the non-injected hemisphere (Fig. 3D, H). Immunohistochemical staining confirmed that NT shRNA control cells in the tumor core had higher expression of hepaCAM as compared to cells expressing HEPACAM shRNAs (Supplemental Fig. 9).

**Figure 3.**
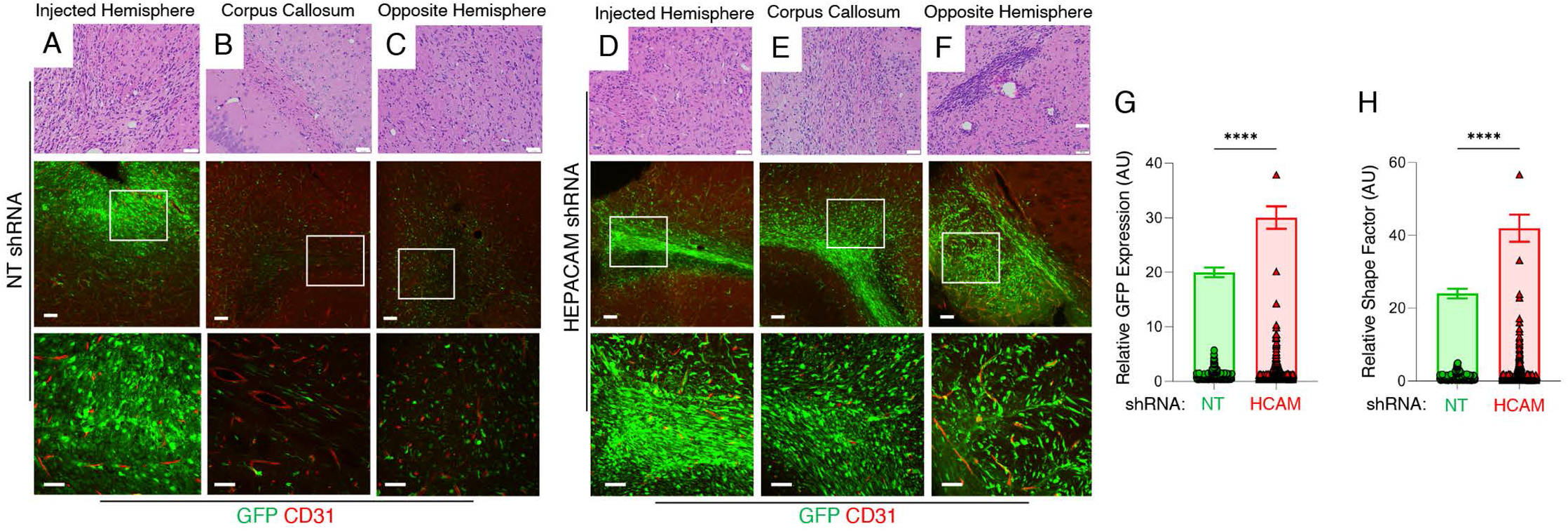
HEPACAM suppresses GSC invasion in the brain. **(A-F);** Coronal sections through the striatum of mice harboring xenograft tumors formed from human GSC6-27 cells expressing NT shRNAs (A-C) or HEPACAM shRNAs (D-F) were H&E-stained (upper panels) or immunofluorescently labeled with anti-GFP and anti-CD31 antibodies (middle and lower panels) to visualize GBM cells and blood vessels, respectively. Shown are images of the injected hemisphere (A, D), the corpus callosum (B, E) and the opposite (contralateral) hemisphere (C, F). Note that in comparison to tumors derived from GSCs expressing NT shRNAs, cells expressing HEPACAM shRNAs showed enhanced invasion through the corpus callosum and distal growth in the opposite hemisphere. The boxed regions in the middle panels are shown at higher magnification in the lower panels. Scale bars: top panel, 50 μm, middle panel, 100 μm and lower panel, 20 μm. **(G, H);** Quantitation of GSC6-27 cell invasive growth, revealing HEPACAM-dependent increases in cell invasion through the corpus callosum and colonization of the contralateral hemisphere in comparison to cells expressing control NT shRNAs. Note that enhanced GFP expression in the corpus callosum confirms the presence of higher number of invasive GSCs with reduced HEPACAM levels compared to GSCs expressing NT shRNAs. These invasive GSCs displayed marked changes in cellular shape (Student’s t-test for comparison, n>100 cells), ****p<0.0001.

Reactive glial cells, and in particular astrocytes and microglia, are significant components of the GBM microenvironment (22). Therefore, spatial localization of microglia and astroglia were next analyzed in xenograft tumors by immunofluorescence using Iba1 and GFAP antibodies, respectively. There were increased numbers of Iba1^+^ microglial cells in HEPACAM shRNA contralateral tumors, as revealed by double labeling with anti-Iba1 and anti-CD31 antibodies (Fig. 4A-G). Similarly, we detected elevated numbers of GFAP^+^ astrocytes in contralateral lesions expressing HEPACAM shRNAs in comparison to control tumors (Fig. 4H-N).

**Figure 4.**
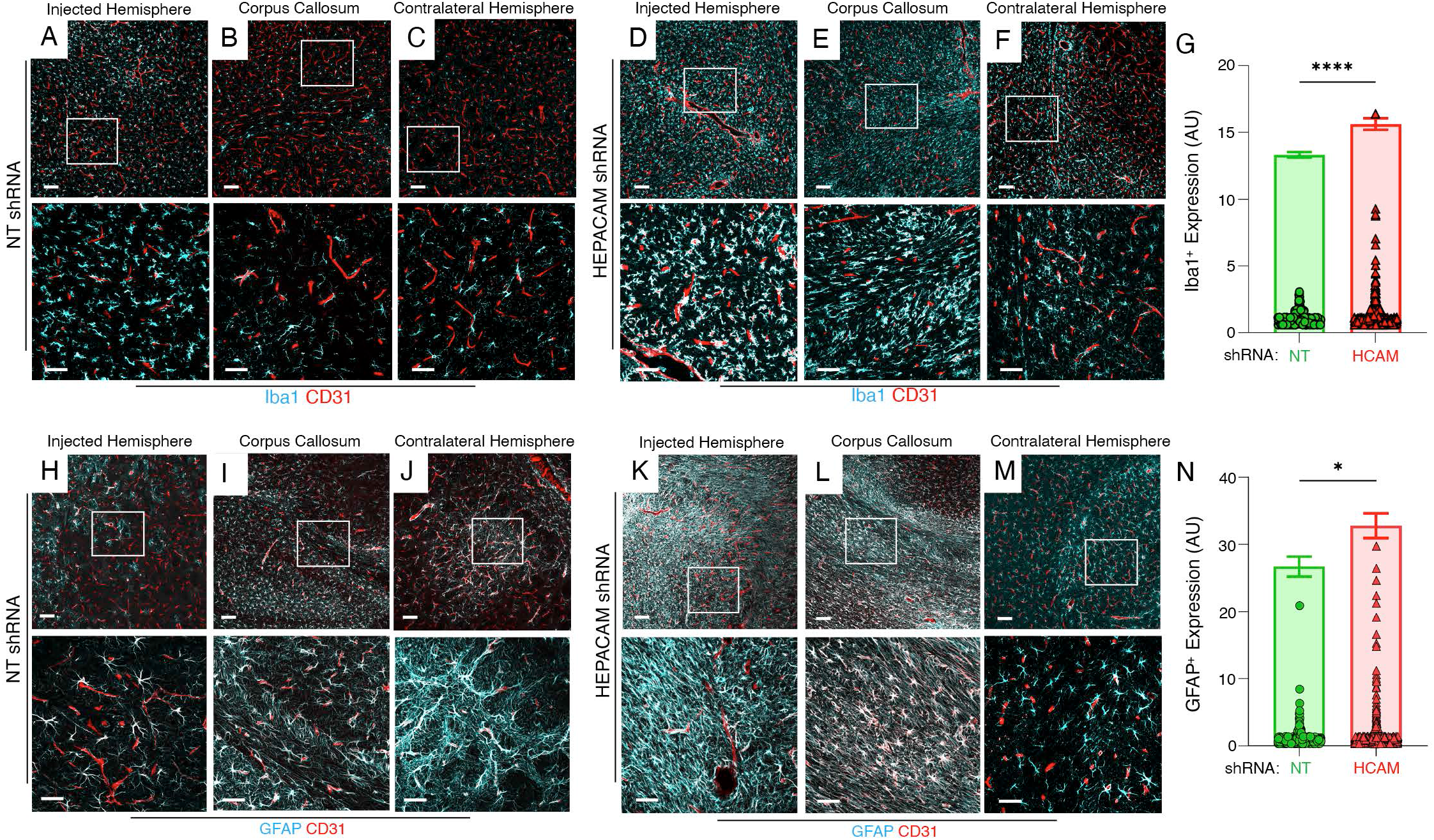
HEPACAM-dependent activation of astrocytes and microglia in the GBM microenvironment. **(A-F);** GSCs expressing NT control shRNAs (A-C) or HEPACAM shRNAs (D-F) were analyzed by double immunofluorescence using anti-CD31 (red) antibody to detect vascular endothelial cells combined with anti-Iba1 (cyan) antibody to detect microglial cells in the injected hemisphere (A, D), corpus callosum (B, E), or contralateral hemisphere (C, F). Note that in comparison to blood vessels in control tumors, the HEPACAM shRNA tumors have increased numbers of Iba1^+^ reactive microglia. The boxed areas in the upper panels are shown at higher magnification in the lower panels. Scale bars: top panel, 100 μm and middle panel, 200 μm. **(G);** Quantitation of increases in Iba1^+^ microglia in HEPACAM shRNA tumors in comparison to NT shRNA control tumors, ****p<0.0001. **(H-M);** GSCs expressing NT control shRNAs (H-J) or HEPACAM shRNAs (K-M) were analyzed by double immunofluorescence using anti-CD31 (red) antibody to detect vascular endothelial cells combined with anti-GFAP (cyan) antibody to detect astrocytes in the injected hemisphere (H, K), corpus callosum (I, L), or contralateral hemisphere (J, M). Note that in comparison to blood vessels in control tumors, the HEPACAM shRNA tumors have increased numbers of GFAP^+^ reactive astrocytes. The boxed areas in the upper panels are shown at higher magnification in the lower panels. Scale bars: top panel, 100 μm and lower panel, 200 μm. **(N);** Quantitation of increases in GFAP^+^ astrocytes in HEPACAM shRNA tumors in comparison to NT shRNA control tumors, *p<0.05.

### HepaCAM suppresses focal adhesion signaling and interactions with ECM

To identify relevant hepaCAM-mediated signaling functions in GSCs we utilized reverse-phase protein arrays (RPPA), which are antibody-based high-throughput systems to study protein signaling cascades (23). Detergent-soluble lysates from GSC6-27 cells expressing HEPACAM shRNAs or control NT shRNAs (n=3 per cell type) were tested. Several proteins involved in focal adhesion signaling were differentially expressed and/or phosphorylated in GSC6-27 cells expressing HEPACAM shRNAs (Fig. 5A). These include Src at the activating tyrosine 416 (Y416) with a concomitant decrease in phosphorylation along with enhanced tyrosine phosphorylation (Y925) of focal adhesion kinase (Fig. 5A). Next, immunoblot experiments were performed to validate elevated levels of focal adhesion protein expression/phosphorylation in GSCs expressing HEPACAM shRNAs. Antibodies recognizing the phosphorylated focal adhesion proteins paxillin and p130Cas, which are substrates for Src, revealed elevated levels in cells expressing HEPACAM shRNAs (Fig. 5B). The cell surface protein β1 integrin is also upregulated in GSC6-27 cells expressing HEPACAM shRNAs (Fig. 5B). Given the hepaCAM-dependent differences in focal adhesion protein phosphorylation, as well as increased expression of β1 integrin, we next analyzed hepaCAM-dependent GSC adhesion to the ECM. GSC6-27 cells expressing control NT shRNAs or HEPACAM shRNAs were plated on various ECM proteins including fibronectin, laminin, collagens and fibrinogen. Reduced hepaCAM expression lead to increased adhesion to fibronectin (Fig. 5C) as well as enhanced cell-spreading and higher levels of paxillin in focal adhesions (Fig. 5D-F).

**Figure 5.**
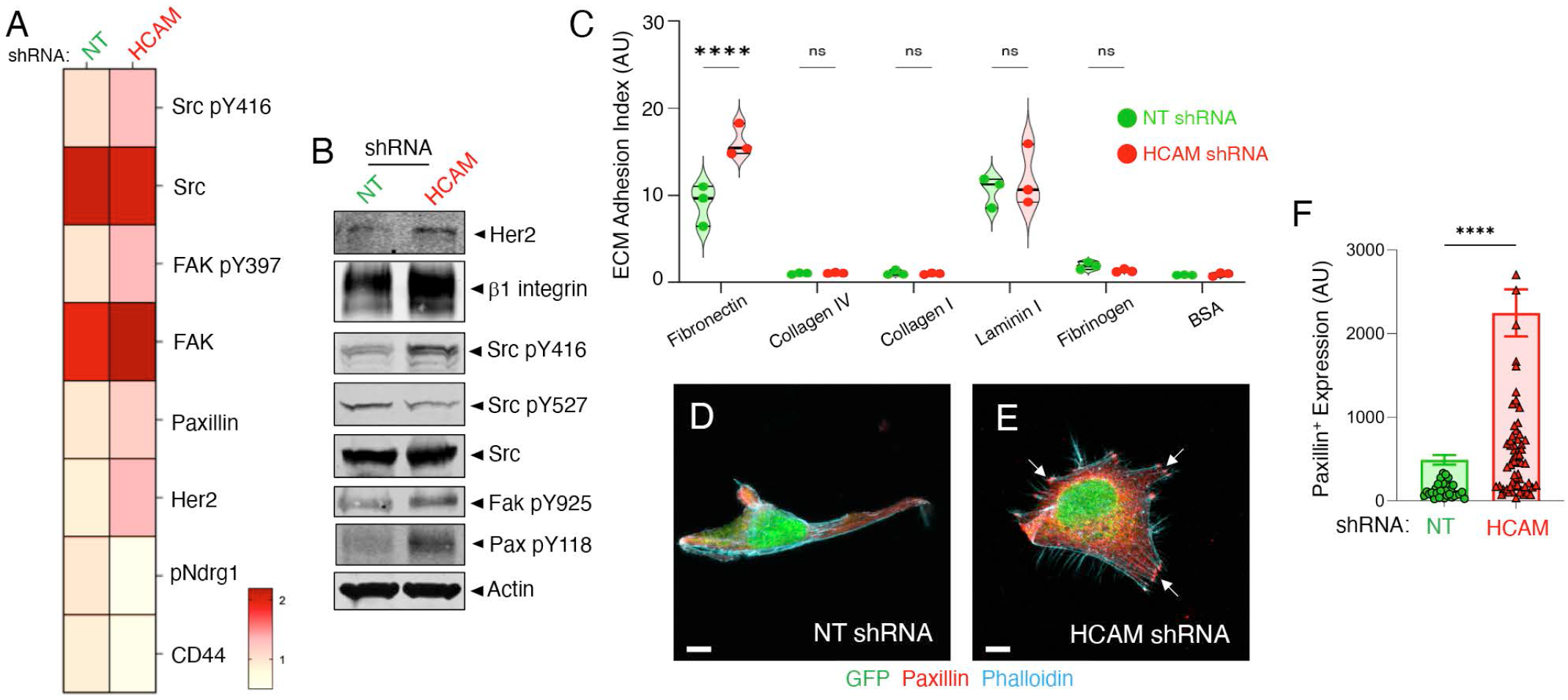
HepaCAM controls focal adhesion signaling dynamics in GSCs. **(A);** RPPA heat map summarizing select proteins that show statistically significant differences in expression and/or phosphorylation in GBM cells expressing HEPACAM shRNAs versus control NT shRNA cells. Proteins displaying reduced (white) or elevated (red) expression and/or phosphorylation are shown for HEPACAM shRNA cells versus NT control cells. **(B);** Immunoblots of detergent-soluble lysates from GSC6-27 cells expressing HEPACAM shRNAs versus control NT shRNAs shows differential expression and/or phosphorylation of select adhesion and signaling proteins identified by RPPA (A). **(C);** GSC6-27 cells expressing control NT shRNAs or HEPACAM shRNAs were added to tissue culture wells coated with the indicated ECM proteins and cell adhesion was quantified. Note that reduced HEPACAM expression leads to significantly increased cell-ECM adhesion selectively to fibronectin. Differences in cell adhesion to other ECM proteins are not statistically significant. (ANOVA and Tukey post-hoc analysis for comparisons, n=3, ****p<0.0001). **(D, E);** GFP^+^ GSC6-27 cells expressing control NT shRNAs (D) or HEPACAM shRNAs (E) plated on fibronectin were labeled with phalloidin (cyan) to visualize the F-actin network in combination with paxillin (red) to identify focal adhesions. Note that in comparison to control GSCs, cells expressing low levels of hepaCAM have increased Factin protrusions and more prominent focal adhesions (arrows). **(F);** Quantitation of focal adhesions in GSC6-27 cells expressing control NT shRNAs or HEPACAM shRNAs. Note the higher expression of the focal adhesion protein paxillin. (Student’s t-test for comparison, n>10 cells per shRNA type, ****p<0.0001).

### Identification of HEPACAM-regulated genes in GSCs

We next performed quantitative RNA sequencing (RNAseq) on human GSCs expressing NT shRNAs or shRNAs targeting HEPACAM (n=3 per cell type) (Fig. 6A). Bioinformatic filtering (FDR 0.05, and fold change > 2 or < 2) reveals that 327 genes are differentially expressed in a HEPACAM-dependent manner, with the differentially expressed genes in GSCs expressing control NT shRNAs versus HEPACAM shRNAs visualized in heat map (Fig. 6B) and volcano plot (Fig. 6C) formats. Gene set enrichment analysis revealed several pathways linked to HEPACAM, including various metabolic pathways as well as pathways linked to focal adhesion signaling, cell-cell adherens junctions, and actin cytoskeletal dynamics (Fig. 6D). Collectively, these various in vitro and in vivo data reveal that hepaCAM is critical for suppressing cell invasion via regulation of focal adhesion signaling pathways and gene expression events in GBM (Fig. 7). High levels of hepaCAM promote GSC proliferation and self-renewal whereas cells with low levels of hepaCAM are more invasive due to increased ECM adhesion and activation of focal adhesion signaling pathways.

**Figure 6.**
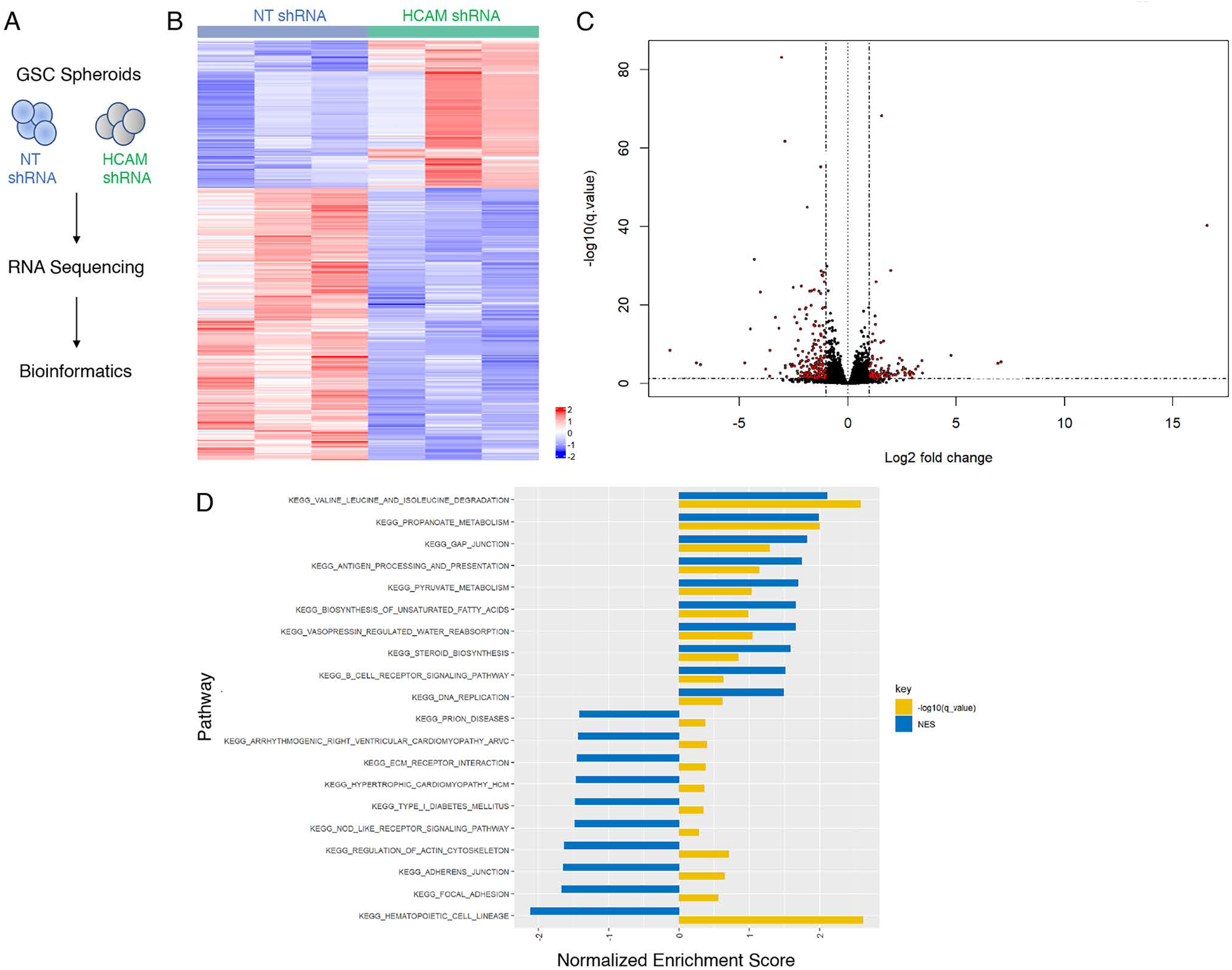
Profiling HEPACAM-regulated gene expression in GSCs. **(A);** Strategy for profiling GSCs to identify genes regulated by MLC1 and/or HEPACAM by quantitative RNAseq. **(B, C);** The 327 differentially expressed genes in GSCs expressing NT shRNAs versus HEPACAM shRNAs shown in heat map (B) and volcano plot (C) formats. Differentially expressed genes were identified using the EdgeR package with adjusted p-value cutoffs <0.05 and log2 fold changes > 2. **(D);** Multiple signaling pathways were identified by gene set enrichment analysis based on differential HEPACAM expression. The normalized enrichment score (NES) and the log transformed (-log) q-values are shown for the top 20 pathways.

**Figure 7.**
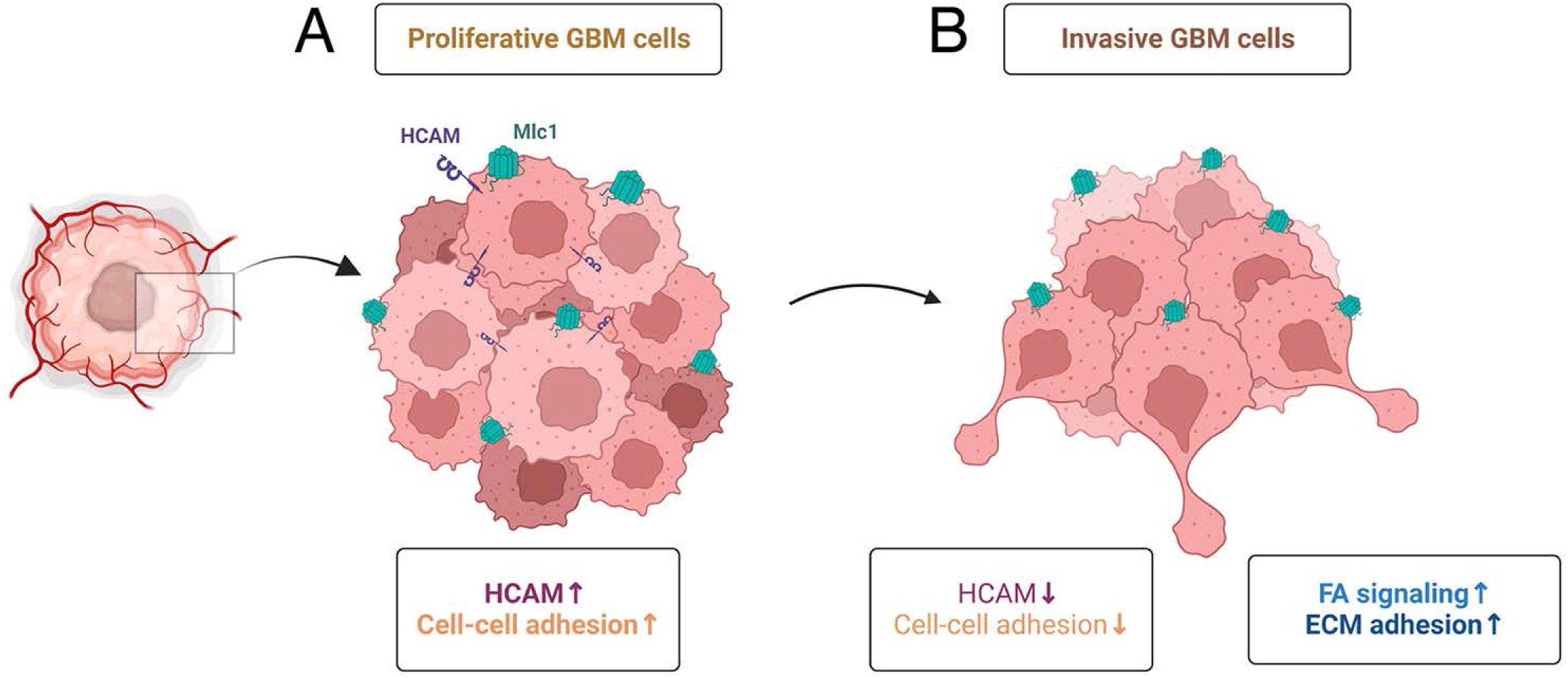
A model for hepaCAM control of GBM cell invasion in the brain microenvironment. **(A);** HepaCAM is expressed in proliferative GBM cells, where it regulates cell-cell adhesion with other GBM cells in the tumor core. **(B);** Reduced hepaCAM expression promotes GBM cell invasion throughout the brain parenchyma. Invasive GBM cells display reduced cell-cell adhesion and enhanced ECM adhesion associated with elevated signaling pathways involving integrins and focal adhesion proteins.

## Discussion

A central conclusion from this study is that hepaCAM is involved in controlling GSC invasion versus growth through balancing focal adhesion signaling dynamics. HepaCAM promotes GSC proliferation and self-renewal by maintaining cell-cell adhesion, Intercellular adhesion also limits dispersal from the main tumor mass. GBM cells with low hepaCAM levels have reduced cell-cell adhesion and increased cell-ECM adhesion, which facilitates invasion through the brain parenchyma. One additional function for hepaCAM may be to suppress Mlc1, with loss of hepaCAM expression leading to Mlc1 activation and enhanced GBM cell invasion. A prior report has linked MLC1 to EGFR activation in cultured astrocytes (24). MLC1 has also been linked to EGFR signaling in low-grade human gliomas harboring R132H mutations in the IDH1 gene Hence, it is enticing to speculate the loss of HEPACAM in GBM cells causes an imbalance in RTK expression and/or signaling via Mlc1. However, the RPPA analyses identify connections between hepaCAM, integrins, and focal adhesion components, which do not overlap with RPPA pathways regulated by Mlc1 in GSCs (10), suggesting that hepaCAM functions at least partially independently of Mlc1 in GSCs.

In most adult tissues and organs hepaCAM is expressed at very low levels, with loss of expression correlated with cell transformation. Indeed, studies of epithelial cancers of the prostate, lung, and colon have shown that HEPACAM has tumor suppressor-like functions, with diminished HEPACAM expression leading to enhanced cell proliferation and metastasis (25–28). These data are different than our results with GBM, which includes GSCs, indicating that hepaCAM functions are influenced by cellular context, oncogenic gene mutations, and possibly other factors in the tumor microenvironment. It will be important to determine the regulatory mechanisms that control HEPACAM gene expression in different GBM cell populations and understand how these events are regulated during tumor progression and recurrence after therapy. A prior report in U373 human glioma cells has shown that forced expression of hepaCAM suppresses growth and induces differentiation (29). Unlike GSCs, we have found that adherent GBM cell lines such as U373 and U87 express very low or undetectable levels of endogenous hepaCAM as determined by analysis of mRNA levels in the NCBI GEO database and immunoblotting GBM cell lysates (A.D., J.M.L., and J.H.M., unpublished data). These results call into question the pathophysiological relevance for exogenous expression of hepaCAM and its impact on highly passaged glioma cell lines. However, high levels of GBM cell invasion do correlate with more stem-like properties (30), suggesting that hepaCAM inhibition may affect GSC differentiation. Hence, it will be important to investigate if hepaCAM impacts expression of markers linked to GSC differentiation status and/or lineage specification.

In GSCs manipulated to express low levels of hepaCAM we detect increased focal adhesion signaling via Src and FAK, higher β1 integrin expression, and enhanced cell spreading on ECM, suggesting that hepaCAM normally suppresses integrin activation and signaling. The cytoplasmic domain of integrin β subunits is involved in the recruitment of proteins, including talins and kindlins, that promote inside-out integrin activation (31). HepaCAM contains a ~125 amino acid cytoplasmic tail with largely unknown functions (1,32). It is possible that intracellular signaling effectors that bind to hepaCAM suppress inside-out integrin activation and dampen ECM adhesion in GBM cells. Alternatively, the extracellular IgG domains of hepaCAM may promote interactions with integrins to dampen adhesion to the ECM. Interestingly, GSCs with low levels of hepaCAM show higher adhesion to the ECM protein fibronectin, which is a molecular marker for the aggressive mesenchymal GBM sub-type (33). We have observed hepaCAM-dependent shape alterations of cells invading across the corpus callosum in the xenograft models, indicating possible links between HEPACAM and the mesenchymal sub-type of invasive GBM cells. The major cell surface receptor for fibronectin is α5β1 integrin (34). A prior report has shown that targeting β1 integrins increases the efficacy of anti-angiogenic therapies in GBM (35) with more specific targeting of α5β1 integrin blocking invasive growth and inducing apoptosis (36). We are currently determining if the hepaCAM extracellular IgG domains or intracellular signaling tail differentially regulate α5β1 integrin-ECM affinity and/or focal adhesion signaling functions. The lipid raft protein caveolin-1/Cav1 couples β1 integrin to the tyrosine kinase Fyn (37) to activate the Ras-Erk pathway (38). Cav1 can also activate α5β1 integrin in GBM cells, which stimulates invasive growth (39). It will be interesting to determine if hepaCAM levels in GBM cells control the balance between α5β1 integrin signaling via Cav1/Fyn to promote proliferation versus invasion via activated α5β1 integrin in focal adhesions.

One gene that is upregulated in GSCs when HEPACAM is silenced is AQP4, which encodes the water channel aquaporin 4/Aqp4. In astrocytes, Aqp4 is a component of the DGC where it regulates water transport at the BBB (9). HepaCAM is also a component of the DGC and patients with MLC disease develop brain edema due to defective water homeostasis at the BBB (40). Upregulation of AQP4 in GBM is linked to increased tumor cell invasion (41), which supports functional connections to hepaCAM. Regulation of water influx mediated by aquaporins at the leading edge is critical for cell shape changes during polarization and invasion (42). Collectively, these data suggest connections between hepaCAM, aquaporin-4 and focal adhesion dynamics in invasive GBM cells.

A recent report has shown that hepaCAM is essential for the establishment of astrocyte territories in the brain, with genetic deletion of Hepacam leading to aberrant astrocyte positioning due, in part, to connexin-mediated cell-cell communication (5). Our data showing that reduced hepaCAM protein levels in GBM cells correlate with invasive behaviors suggest that spatial location of GBM cells in a tumor may also be regulated by hepaCAM. The data from the xenograft models also indicate that GBM cell communication with tumor-associated astrocytes may be dependent, in part, on intercellular adhesion via hepaCAM. GBM cells expressing low levels of hepaCAM may have defective contact communication with astrocytes. The roles for astrocytes in the GBM microenvironment remain uncertain, with some publications supporting tumor promoting functions whereas other studies show tumor suppressive roles (43). It will be important to determine if levels of hepaCAM in GBM cells control interactions with different sub-populations of astrocytes or other glial cell types such as microglia that impact tumor malignancy. HepaCAM also interacts with aquaporin 4/AQP4, Trpv4 and other components of the DGC in astrocytes, which also includes the intracellular adapter proteins α-dystrobrevin and dystrophin, α/β-dystroglycan transmembrane proteins, and ECM proteins such as agrin and laminin (9). The DGC complex is enriched at the astrocyte–blood vessel interface and has important but understudied roles in regulating BBB development and integrity (44). Functions for hepaCAM in the DGC of GBM cells remains largely unknown and will be an important area of future investigation.

Anti-vascular therapies that target angiogenic blood vessels in GBM, including the cyclic RGD peptide mimetic Cilengitide that blocks ECM adhesion by the endothelial cell-expressed integrins αvβ3 and αvβ5 have largely failed in GBM clinical trials (45). In addition, anti-VEGF agents such as the neutralizing antibody bevacizumab or small molecule inhibitors of VEGF receptor tyrosine kinases, do not improve overall patient survival (46,47). Indeed, in pre-clinical mouse models of brain cancer and in nearly 50% of patients with GBM (48–50), bevacizumab treatment leads to a pathological burst in tumor cell invasion (51). Therefore, it is enticing to speculate that down-regulation of HEPACAM may be one additional mechanism used by GBM cells to diminish adhesion/communication with blood vessels, exit the primary tumor mass and disperse throughout the brain in response to anti-angiogenic therapies. If so, these results would suggest that hepaCAM functions are linked to VEGF receptor signaling pathways, possibly in conjunction with integrins.

## Materials and Methods

### Human GBM cells

Approval for the use of human specimens was obtained from the Institutional Review Board (IRB) at the University of Texas MD Anderson Cancer Center. The IRB waived the requirement for informed consent for previously collected residual tissues from surgical procedures stripped of unique patient identifiers according to the Declaration of Helsinki guidelines. Glioblastoma stem cells (GSCs); GSC6-27. GSC2-31 and GSC214 were cultured from freshly resected human tumors [20] and, grown and maintained in complete GSC media; DMEM Ham’s F12 50/50 medium containing L-glutamine (Corning, 10-090-CV), supplemented with 1X B27 supplement (Life Technologies, 17504-044), 20 ng/mL EGF (Biosource, PHG0313), 20 ng/mL bFGF (Biosource, PHG0021), and 1X Penicillin-Streptomycin-antimycotic solution (Corning, MT30004CI). When GSCs developed neurosphere-like spheroids in culture, they were passaged by dissociation using 50 μL Accutase™ (Innovative Cell Technologies Inc., AT104) per 1 × 10^6^ cells and maintained in complete GSC media. GSCs were made adherent by withdrawing EGF/FGF from the growth medium and culturing on glass slips coated with poly-L-ornithine (1:100; Sigma, P4957), fibronectin (1:60, Sigma, F0895) and laminin (1:300, Sigma, L2020). Genomic validation of GSCs was performed by DNA short tandem repeat profiling in a CCSG-funded Characterized Cell Line Core Facility. GSCs were routinely tested for mycoplasma using commercially available kits (Thermo Fisher), and only those cells deemed mycoplasma free were used for experiments.

GSCs were infected overnight with concentrated pGIPZ lentivirus at a multiplicity of infection of 1.0. The following clones were used for HEPACAM shRNA: V3LHS_413349, V3LHS_413351, and V3LHS_413352 versus control pGIPZ containing RFP (GE Dharmacon). HEK293T were transfected with HEPACAM ORF using the Precision LentiORF (pLOC) vector versus control pLOC (GE Dharmacon). All HEK293T were transfected using Lipofectamine 3000 reagent (Thermo Fisher).

### ECM invasion assays

CytoSelectTM 24-well cell invasion assay, basement membrane (Cell Biolabs, CBA-110) was used for in vitro cell invasion assay (colorimetric) of NT shRNA (n=3) and HEPACAM shRNA (n=3) GSCs by following manufacturer’s instructions. Spheroids were dissociated as per previously described protocol. ECM coated cell culture inserts (8 μm pore size) were seeded with 1 × 10^5^ viable GSCs in complete GSC medium and the lower chambers were filled with chemoattractant; 10% FBS (Sigma, F0926) containing DMEM Ham’s F12 50/50 medium, supplemented with 1X antibiotic-antimycotic solution, followed by incubation at 37°C and 5% CO_2_ for 24 hours. Invasive cells degraded the ECM proteins and passed through the pores of the membrane of the cell culture insert while non-invading cells remained inside the insert and were removed by a cotton swab. Invasive cells adhering to the outer side of the inserts were stained and quantified by measuring OD (absorbance) at 560 nm (BioTek, Synergy HTX multimode reader).

### RNA sequencing and bioinformatics

Spheroids from GSC6-27 NT shRNA (n=3) and GSC6-27 HEPACAM shRNA (n=3) cultures were washed in cold 1X PBS and total RNA was extracted using RNeasy Plus Mini Kit (Qiagen, 74134) following manufacturer’s guidelines. Following RNA quality validation (RIN≥7), samples were sequenced using NextSeq500 high output 75nt PE flow cell instrument (ATGC Illumina Next Generation Sequencing Core facility) by generating 44 million paired-end reads for each sample. Sequenced reads fastq files were aligned with Star/2.6.0b, number of reads per gene were counted with HTSeq/0.11.0 and the read counts were normalized with “DESeq2”. Plots, hierarchical clustering and Principal Component Analysis (PCA) were used for quality assessment, and normalized log2 read counts were fitted with negative binomial GLM and tested by Wald statistics. Regularized log transformed data was used for visualization, clustering and PCA analysis. Beta-uniform mixture (BUM) model was used to for determining p-values from Wald statistical tests. We analyzed an average of ~45 million paired-end reads generated for each of the six GSC samples (n=3 NT shRNA versus n=3 HEPACAM shRNA). Sequenced reads fastq files were mapped to human reference genomes using Tophat (52). Raw reads were calculated using HTseq (53). Differentially expressed genes, defined as those having logFC > 2 and FDR< 0.01, were obtained with EdgeR package analyses (54). The signal-to noise metric was used to calculate the gene expression differences between cell samples. KEGG pathways were compiled from MsigDB (55). Unsupervised gene set enrichment analysis for all the KEGG pathways was performed using GSEA. Normalized enrichment scores and false discovery rate (FDR) values were calculated under a 1000-fold permutation.

### Pre-clinical mouse models of GBM

Mouse studies were reviewed and approved prior to experimentation under the guidelines of the Institutional Animal Care and Use Committee and The University of Texas M.D. Anderson Subcommittee on Animal Studies, both AAALAC accredited institutions. Male nude (NCR nu/nu) mice were purchased from Jackson Laboratories and used for all GSC implantation experiments. Randomization was not used since all mice used were the same age and sex. Seven-week-old mice were anesthetized and a Hamilton syringe was used to dispense 1.5 × 10^5^ GSC6-27 cells infected with pGIPZ with NT shRNAs or HEPACAM shRNAs (n=10 mice per cell type). The sample size was selected to ensure power analyses using a = 0.05 and an effect size = 0.4 for comparing the two groups using one-way analysis of variance (ANOVA). Mice were euthanized at 16 weeks post injection, perfused with 4% PFA/PBS and brains were sectioned for experimental analyses. Animals were excluded from the analysis if death occurred before 16 weeks after injection.

### Reverse-phase protein arrays

GSCs (+/− HEPACAM shRNAs) were washed twice in ice-cold PBS, then lysed in RIPA or RPPA lysis buffer containing 1%Triton X-100, 50mM Hepes pH 7.4, 150mM NaCl, 1.5mM MgCl_2_, 1mM EGTA, 100mM NaF, 10 mM sodium pyruvate, 1 mM Na_3_VO_4_, 10% glycerol, and a cocktail of protease and phosphatase inhibitors (Roche Diagnostics, Mannheim, Germany) for 20–30 min with frequent mixing on ice, then were centrifuged at 14,000 RPM at 4°C for 15 min to isolate the detergent soluble protein supernatant. The protein concentration was determined using the BCA Assay (Thermo Scientific, Pierce™ BCA, 2327). The optimal protein concentration of lysates for RPPA is about 1.2 μg/μl (1.2 mg/ml). Lysates were denatured in 4% SDS/2-ME sample buffer (35% glycerol, 8% SDS, 0.25M Tris-HCl, pH 6.8; no β-mercaptoethanol) for 5 min at 95°C. Lysates were stored at −80°C and subsequently analyzed in the RPPA core facility at MD Anderson Cancer Center. Samples were serially diluted and probed with 466 antibodies and arrayed on nitrocellulose-coated slides.

### ECM adhesion assays

Cell adhesion assays were performed using ECM cell adhesion array kit (Cell Biolabs, Inc., CytoSelect™ 48-Well Cell Adhesion Assay, CBA-070),) following the manufacturer’s instructions. Spheres were dissociated using Accutase Accutase™ (Innovative Cell Technologies Inc., AT104) followed by centrifugation (1000 rpm, 5 mins., RT). Viable cell count was obtained (Beckman Coulter, Vi-cell Analyzer) and cells were gently re-suspended in 4 mls of assay buffer. Subsequently, 150 μl of the cell suspension containing 1. ×10^6^ viable GSCs in serum free media (GSC media lacking growth supplements) was added to each well and incubated for 90 minutes at 37°C and 5% CO_2_. The non-adherent cells were gently removed by rinsing each well four times with 1X PBS. The resulting adherent cells were fixed and stained at RT for 10 min. Cells were washed four times with deionized water. Stain was extracted from each well and absorbance readings were taken at 560 nm (BioTek, Synergy HTX multi-mode reader).

### Immunofluorescence

Fixed brain samples were embedded in 4% agarose and sectioned at 100 μm on a vibratome and stored in 1X PBS at 4°C. Alternatively, brains were processed for paraffin embedding and sectioning. Sections were permeabilized and blocked with 10% donkey serum in PBS-T (1X PBS supplemented with 0.1-0.25% Triton-X100) for 1 hour at room temperature (RT), followed by an overnight 4°C incubation with primary unconjugated antibodies diluted in the blocking solution. Immunofluorescence analyses were performed with the following primary antibodies: anti-CD31 (R&D Systems AF3628), GFP (Abcam ab290 or ab13970), GFAP (DAKO Z0334 or Novus NBP1-05198), Iba1 (WAKO 0919741), and Laminin (Sigma L9393). The sections were then washed with PBS-T and incubated with secondary antibodies (1:500 dilution) in the blocking solution for one hour. Sections were again washed 3 times with PBS-T, then briefly washed with PBS. The sections were mounted on pre-treated microscope slides, sealed using Vectashield with DAPI mounting media (Vector Laboratories, Inc)) and kept at 4°C until imaging. Confocal Images were acquired using an Olympus FLUOVIEW FV3000 10X, 20X and 30X objectives. All comparative images were taken with the same laser power and gain settings in order to make qualitative and quantitative comparisons between staining levels in different samples. Multiple fields of view were imaged from biological replicates.

### GBM cell fluorescence staining

Chamber slides (Nunc, Lab-Tek) were coated with 1X PBS containing fibronectin (1:60, Sigma, F0895) overnight at 4°C. Chambers were rinsed thrice with 1X PBS. 400 μL of 1X PBS containing 2% BSA (Hyclone, SH30574) was added to each well and incubated at RT for 1 hour. GSC spheroids in culture were dissociated by Accutase™ (Innovative Cell Technologies Inc., AT104) and single cells were resuspended in DMEM Ham’s F12 50/50 medium containing 2% BSA. Viable cell count was obtained (Beckman Coulter, Vi-cell Analyzer) and 5×10^3^ viable GSCs were added to each well containing BSA-DMEM Ham’s F12 50/50 medium followed by incubation at 37°C and 5% CO_2_ for 2 hours. Adherent GSCs were gently rinsed (5 mins, RT) with 1X PBS, fixed with 4% paraformaldehyde in 1X PBS at RT for 10 mins and rinsed thrice (5 mins each, RT) with ice-cold 1X PBS. Cells were permeabilized with 0.1% Triton-X 100 for 10 mins at RT and rinsed twice with 1X PBS. Cells were blocked with 1% BSA in 1X PBS for 30 mins at RT and incubated (4°C) overnight with primary unconjugated antibodies; anti-Paxillin (1:100, Thermo Fisher, AHO0492), anti-HEPACAM (1:100, R&D Systems, MAB4108) and anti-GFP (1:1000, Abcam, ab13970) diluted in blocking buffer. Cells were rinsed thrice (5 mins each, RT) with wash buffer (0.1% BSA in 1X PBS) and incubated (RT) for 45 mins with fluorochrome conjugated secondary antibodies; anti-chicken-Alexa Fluor 488 (1:500, Jackson Immunoresearch, 703-545-155), anti-mouse-Alexa Fluor 594 (1:500, Jackson Immunoresearch, 715-585-150) and anti-rabbit-Alexa Fluor 594 (1:500, Jackson Immunoresearch, 711-585-152) diluted in blocking buffer. Additionally, Phalloidin-iFluor 647 (1X, Abcam, ab176759) was used (in combination with secondary antibody solutions) to stain actin filaments. Cells were rinsed thrice (5 mins each, RT) with wash buffer, rinsed twice with 1X PBS and finally the chamber slides were sealed using Vectashield with DAPI mounting media (Vector Laboratories, Inc.). Confocal Images were acquired using an Olympus FLUOVIEW FV3000 30X and 60X oil immersion objectives. All images were captured with the same laser power and gain settings in order to make comparisons between staining levels in different samples. Multiple fields of view were imaged from biological replicates.

### Immunohistochemistry

Immunohistochemical staining was performed on formalin-fixed paraffin-embedded mouse brain tissue. The tissue was deparaffinized at 65°C and rehydrated in decreasing concentrations of alcohol. Heat-induced antigen retrieval was performed using a Na-citrate buffer at pH 6. The tissue was blocked in 10% serum of the same species as the secondary antibody for one hour at room temperature. The tissue was then incubated overnight at 4°C in primary unconjugated antibodies diluted in blocking buffer. Immunohistochemical analyses were performed with the following primary antibodies: rabbit anti-hepaCAM (Proteintech 18177-1-ap, 1:200 dilution), anti-Mlc1 antibody (Novus Biologicals, NBP1-81555), human-specific goat anti-Vimentin (R&D Systems, AF2105). Slides were then washed in 1X PBS and incubated with biotinylated secondary antibodies (1:500-1:1000) in the blocking buffer for 1 hour at RT. Following another series of washes in PBS, the slides were incubated in ABC reagent (Vector Laboratories, PK-4000) for 30 minutes at RT. Following more washes with PBS, ImmPACT DAB Substrate (Vector Laboratories, SK-4105) was added for 5-10 minutes before rinsing the slides in ddH_2_O and counterstaining with hematoxylin for 10-15 seconds. After rinsing in cold tap water, the slides were then dehydrated in increasing concentrations of alcohol and mounted using mounting media and stored until images were collected. Light microscope images were acquired using an Olympus BX43 with a 20x objective.

### Immunoblotting

Whole cell lysates were collected under normal culture conditions by lysing spheres in radioimmunoprecipitation assay buffer (RIPA; 50mM Tris-HCl pH 8.0, 150 mM NaCl, 1% sodium deoxycholate, 1% triton X-100, 0.1% SDS, 1% NP-40 and 1 mM EDTA) containing protease and phosphatase inhibitors (A32955, A32957, Thermo Scientific) to obtain soluble protein fractions. Total protein was measured by Pierce™ BCA protein assay kit (Thermo Scientific, 23227), and then denatured at 95°C for 10 min in 4X Laemmeli buffer (Bio-Rad, 1610747) containing 2.5% 2-mercaptoethanol (Sigma, M6250). 30-75 ug of protein was resolved on 4-15%, 10% or 12%Tris-glycine gels. Immunoblotting was performed with nitrocellulose membranes (Bio-Rad, 1620112, 1620115), blocked using Odyssey TBS-based blocking buffer (LI-COR), and then incubated with specific primary antibodies diluted in blocking buffer supplemented with 0.1% Tween-20 overnight at 4°C. The following primary antibodies were used for immunoblotting: anti-hepaCAM (Proteintech 18177-1-ap, 1:500), anti-Src (CST, 2109, 1:1000), anti-phospho-Src-pY416 (CST, 2101, 1:1000), anti-phospho-Src-pY527 (CST, 2105, 1:1000), anti-p130Cas (CST, 13846, 1:1000), anti-phospho-p130Cas-pY410 (CST, 4011, 1:1000), anti-phospho-FAK-pY397 (CST, 8556, 1:1000), anti-phospho-FAK-pY925 (CST, 3284, 1:1000), anti-phospho-FAK-pY576/577 (CST, 3281, 1:1000), anti-phospho-paxillin-pY118 (CST, 2541, 1:1,000), anti-phoshpho-Tyrosine-pY99 (Santa Cruz Biotech, sc-7020, 1:1000), anti-Integrin β1 (Abcam, ab183666, 1:2000), anti-Her2 (CST, 2242, 1:500), anti-phospho-Her2-pY1196 (CST, 6942, 1:1000), anti-phospho-NDRG1-T346 (CST, 3217, 1:1000), anti-NDRG1 (CST, 5196, 1:1000) and anti-CD44 (CST, 3578, 1:1000). Target proteins were normalized to total cellular/housekeeping proteins: anti-α-actinin (Abcam, ab18061, 1:3000), anti-β-actin (Abcam, ab6276, 1:3000) and anti-GAPDH (Sigma, G8796, 1:2000). Blots were incubated with fluorochrome conjugated secondary antibodies (IRDye 800CW goat anti-rabbit and IRDye 680RD goat anti-mouse, at 1:15,000 dilutions) (LI-COR) in the dark at RT for 30 minutes. Dual-channel infrared scan and quantitation of immunoblots were conducted using the Odyssey CLx infrared imaging system with Image Studio (Ver. 5.2) (LI-COR).

### Image acquisition, analysis and quantitation

Immunofluorescence images were acquired using an Olympus Fluoview FV3000 confocal laser scanning microscope. Multidimensional acquisition was conducted using Z stacks with 2.5 μm slicing intervals at a scan rate of 4ms/pixel with a resolution of at least 1024 × 1024 pixels per slice and digitally compiled in FV31S-SW (version 2.4.1.198). Image acquisition parameters, including exposure time, laser power, gain, and voltages, were fixed for each imaging channel. Immunohistochemistry-labeled images were captured using an Olympus BX43 light microscope. Using ImageJ, all images were scaled (μm) as per the objective lens used for acquisition to measure signals and cellular parameters. Acquired image Z-stacks were projected for maximum intensity to include all signals, but the ‘auto-threshold’ module was used to include only cellular signals for quantitation and exclude non-specific background or noise. Fluorescence signal intensity of channels were measured using the standard “color histogram” module. Cell counts, mean fluorescence signals from single cells and cellular peripheries, major and minor axes for defining cell shape were analyzed using the ‘analyze particles’ algorithm (56).

### Statistical analyses

Quantitation of confocal images was performed using ImageJ software (NIH, USA) (57,58). Graph Pad prism 9.0 was used to plot mean values and all data points (n=3 or greater ± SEM per group, unless otherwise indicated) to compare between experimental and control samples and to determine statistical differences by unpaired Student’s t-test and two-way ANOVA (Tukey post-hoc analysis) at 95% confidence intervals (α value 0.05). Statistical p-values less than *0.05, **0.01, ***0.001 and ****0.0001 were considered significant.

## Acknowledgements

We thank the various members of the McCarty laboratory for insightful comments on the manuscript. We are also grateful to Dr. Daniel Wagner and members of his research group at Rice University for assistance with HEPACAM cDNA sub-cloning. We also thank Drs. Sabbir Khan and Sanjay Singh (MD Anderson Cancer Center) for insightful technical advice with some of the laboratory techniques. This work was supported, in part, by grants to J. H. M. from the Cancer Prevention and Research Institute of Texas (RP180220), the National Institutes of Health (R01NS087635, R21NS103841, and P50CA127001), the Brockman Foundation, and the Terry L. Chandler Foundation (TLC^2^) Foundation from the Heart. The following NCI-funded Cancer Center Support Grant (CCSG) Core Facilities were instrumental in data acquisition: the shRNA and ORFeome Core, the Research Histopathology Facility, the Flow Cytometry and Cellular Imaging Facility, and the Sequencing and Microarray Facility.

