## Supplemental Figures and Legends for "HepaCAM Suppresses Glioblastoma Stem Cell Invasion in the Brain"

#### Supplemental Figure Legends

**Supplemental Figure 1. Analysis of hepaCAM protein expression in GBM and validation of anti-hepaCAM antibody specificity. (A-E);** Anti-hepaCAM immunohistochemical staining of five different human GBM samples. **(F);** A human GBM section was immunohistochemically labeled with species-matched control IgG. Scale bars, 50 μm. **(G);** Lysates from HEK-293T cells transfected with human pLOC lentivirus expressing HEPACAM were mock treated or treated with PNGase to remove N-glycosylation sites. Lysates were then resolved by SDS-PAGE and immunoblotted with anti-hepaCAM antibody. Note the shift in molecular weight for hepaCAM following PNGase treatment.

**Supplemental Figure 2. Identification of genes co-expressed with HEPACAM in the non-cancerous brain and in GBM. (A, B);** Correlation AnalyzeR software was used to identify genes that are co-expressed with HEPACAM in non-cancerous human brain samples (A) or human GBM samples (B). Gene expression data from the ARCHS4 platform were used for analysis.

**Supplemental Figure 3. HEPACAM is predominantly expressed in astrocytes in the mouse and human brain. (A, B);** Analysis of two different open source murine transcriptomic databases reveals that Hepacam mRNA is expressed in astrocytes and oligodendrocytes of the neonatal mouse brain (A) (brainrnaseq.org) and the adult mouse brain (B) (betsholtzlab.org/VascularSingleCells/database.html). **(C);** HEPACAM mRNA is enriched in astrocytes and oligodendrocytes of the adult human brain (twc-stanford.shinyapps.io/human_bbb).The expression data were graphed as fragments per kb of transcript per million (FKPM).

**Supplemental Figure 4. Expression analysis of Hepacam mRNA in different mouse brain regions. (A);** Analysis of the Human Protein Atlas (www.proteinatlas.org) reveals that Hepacam mRNA is predominantly expressed in the post-natal mouse brain. Note the variable levels of expression in different regions of the brain and olfactory bulbs. **(B);** Analysis of Hepacam mRNA in the GTex brain database reveals predominant expression in the mouse brain with varying expression levels in different brain regions.

**Supplemental Figure 5. Analysis of HEPACAM mRNA expression levels in different human cancers.** Analysis of the TCGA dataset across various cancer types reveals elevated expression of HEPACAM in gliomas and liver cancers.

**Supplemental Figure 6. HEPACAM expression is down-regulated in human low grade gliomas. (A-E);** Five different low-grade gliomas were immunohistochemically labeled with anti-hepaCAM, revealing protein enrichment in some perivascular tumor cells and more diffuse expression throughout the tumor. Scale bars, 50 $\mu$m. **(F);** HEPACAM mRNA levels are higher in low-grade gliomas (n=529) versus normal brain (n=207), as determined by analysis of the GTex and TCGA databases. (Unpaired Student's t-test for comparisons, ****p<0.0001).

**Supplemental Figure 7. Analysis of HEPACAM expression in primary versus recurrent human GBM.** The Chinese Glioma Genome Atlas (CGGA), which contains RNA sequencing data from a large number of primary (n=651) and recurrent (n=333) glioma samples, was queried for HEPACAM expression levels. In comparison to primary surgical samples, note the significant reduction in HEPACAM mRNA levels in recurrent glioma samples. (Unpaired Student's t-test for comparison, ****p<0.0001).

**Supplemental Figure 8. HEPACAM controls GSC self-renewal and invasion in vitro. (A);** The percentages of newly formed spheres (>20,000 $\mu m^2$) were recorded daily for 7 consecutive days. GSC231 cells expressing HEPACAM shRNAs showed reduced spheroid formation(ANOVA and Tukey post-hoc analysis for comparison, n=4, ****p<0.0001). **(B);** GSC231 cells expressing HEPACAM shRNAs showed enhanced invasion through basement membrane-coated transwells as compared to NT shRNA control pGIPZ-infected cells (Unpaired Student's t-test for comparison, n=3, **p<0.01). **(C);** The numbers and cross-sectional areas of newly formed spheres were recorded daily for 7 consecutive days. In comparison to control GSC2-14 cells, GSC2-14 cells expressing HEPACAM shRNAs showed reduced spheroid formation (ANOVA and Tukey post-hoc analysis for comparison, n=4, ***p<0.001) **(D);** GSC2-14 cells expressing HEPACAM shRNAs showed enhanced invasion through Matrigel-coated transwells as compared to NT shRNA control pGIPZ-infected cells, (Unpaired Student's t-test for comparison, n=3, *p<0.05).

**Supplemental Figure 9. Analysis of hepaCAM expression in xenograft models of GBM. (A-D);** Coronal sections through the striatum of mice harboring tumors formed from human GSC6-27 cells expressing non-targeting (NT) shRNAs (A, B) or HEPACAM shRNAs (C, D) were immunohistochemically stained with anti-hepaCAM (A, C) or human-specific anti-vimentin antibodies (B, D). Vimentin serves as a general marker for human tumor cells. Shown are images of the injected hemisphere in xenograft mice. Note that in comparison to tumors derived from GSC6-27 cells expressing NT shRNAs which express hepaCAM, cells expressing HEPACAM shRNAs showed reduced hepaCAM protein expression. Scale bars, 50 μm.

### Supplemental Figure 1

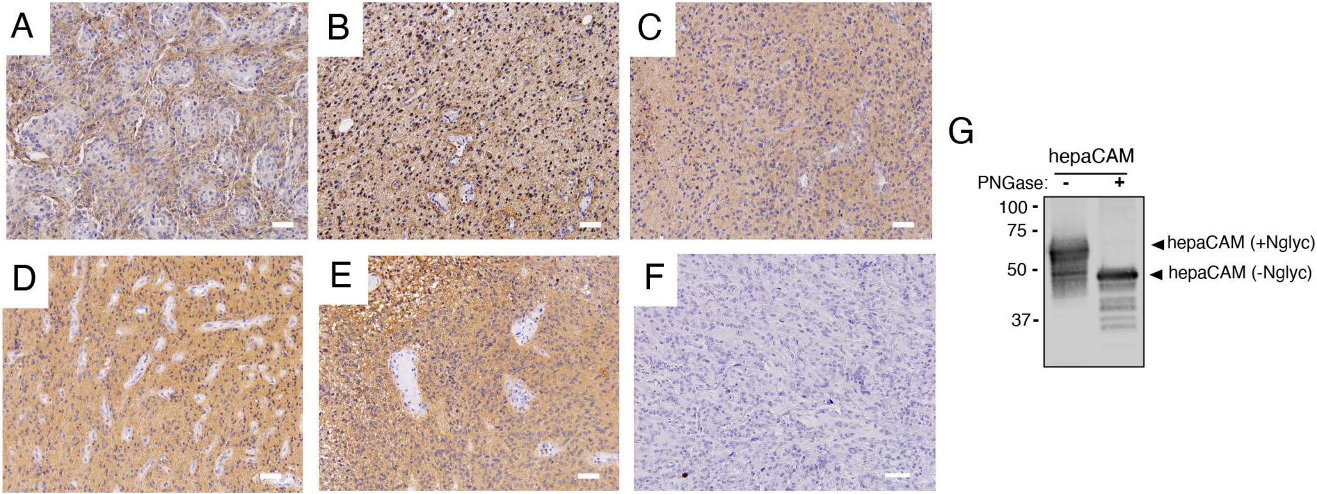

### Supplemental Figure 2

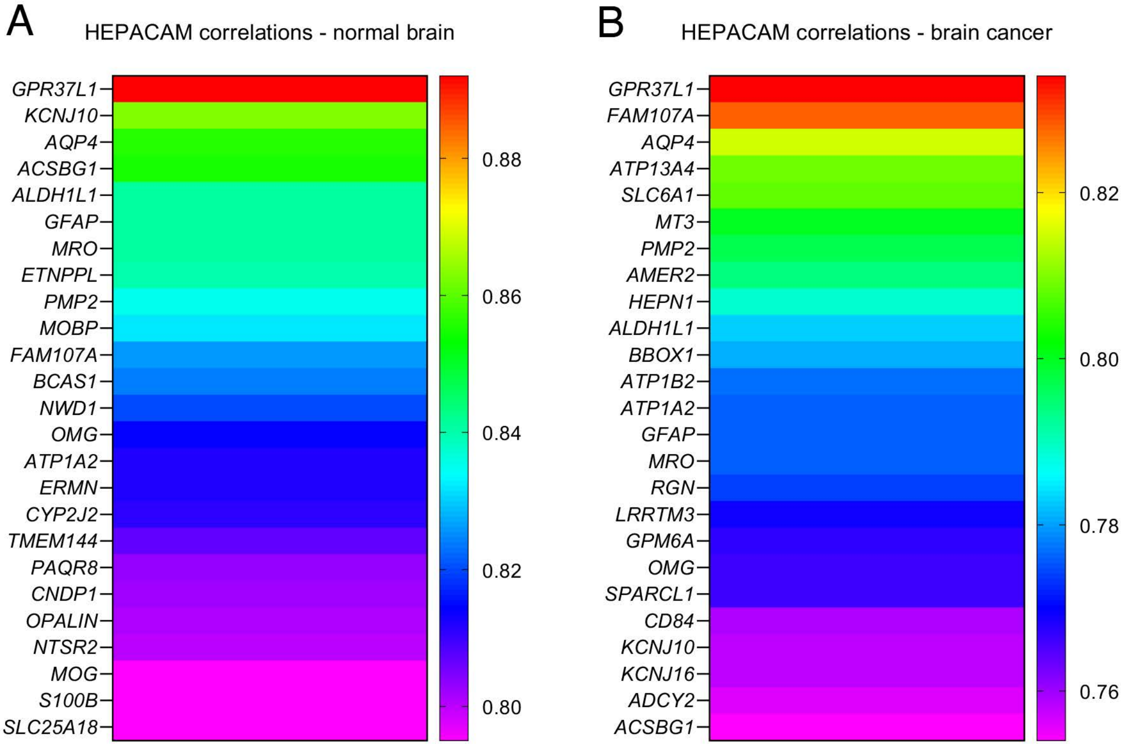

Supplemental Figure 3

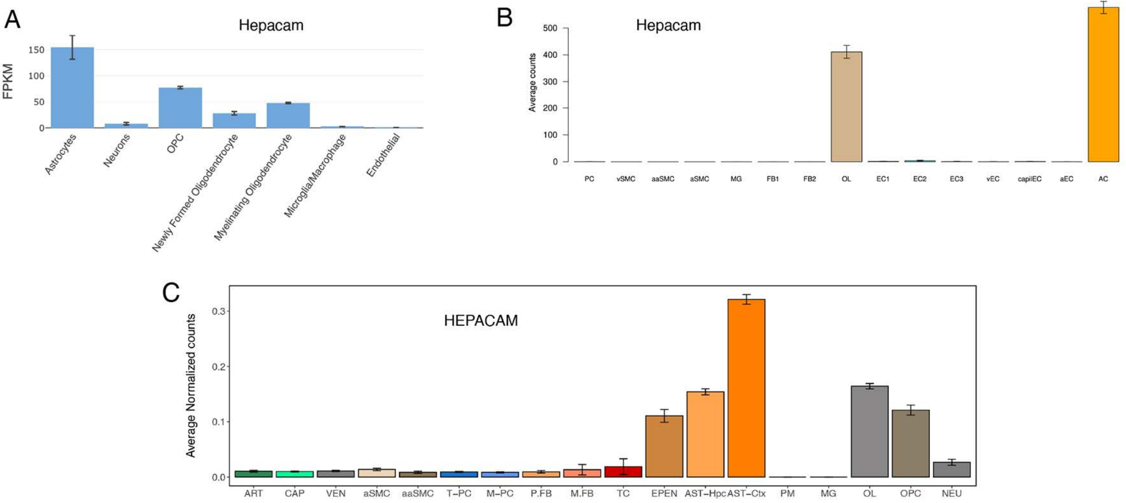

### Supplemental Figure 4

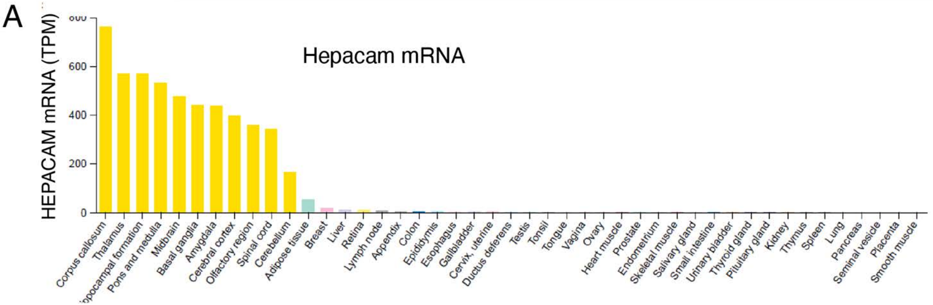


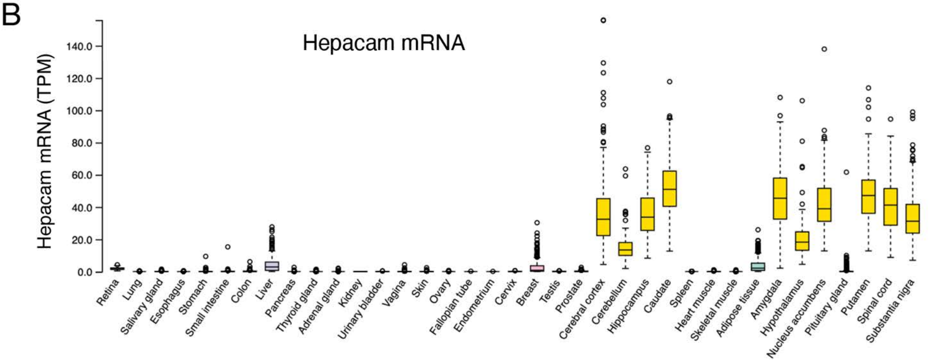

Supplemental Figure 5

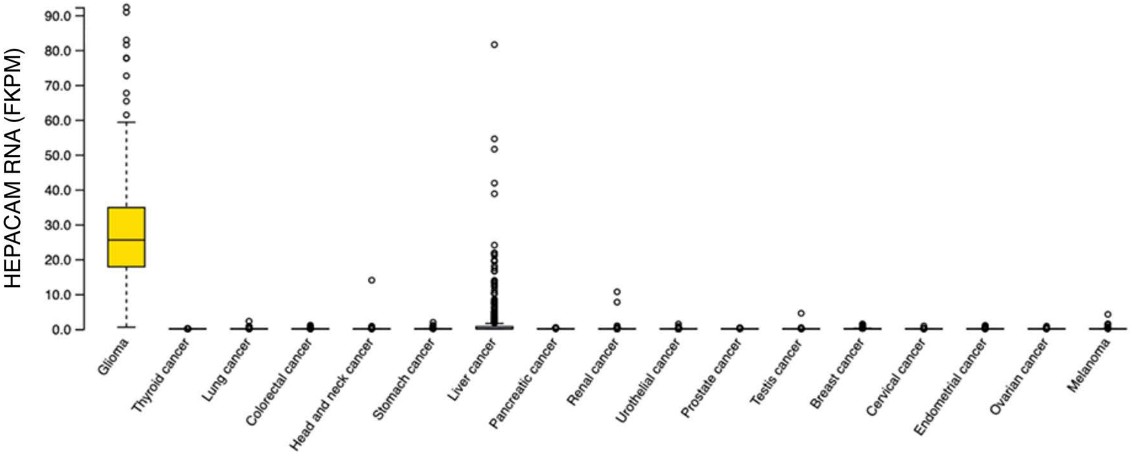

### Supplemental Figure 6

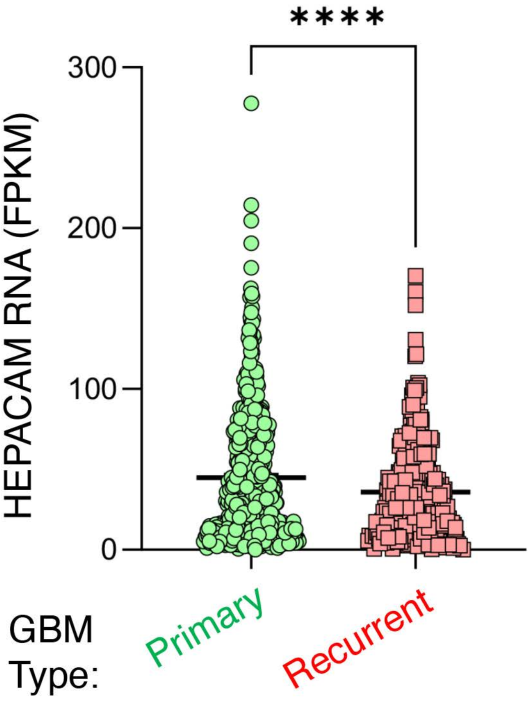

Supplemental Figure 7

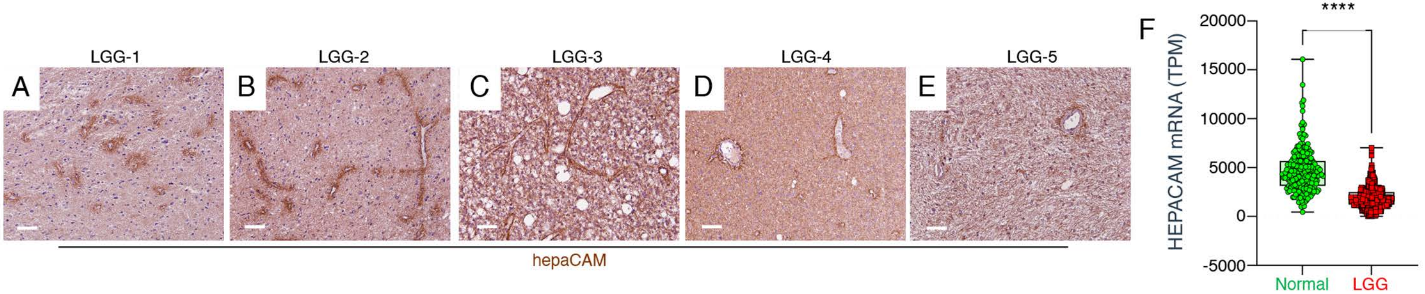

### Supplemental Figure 8

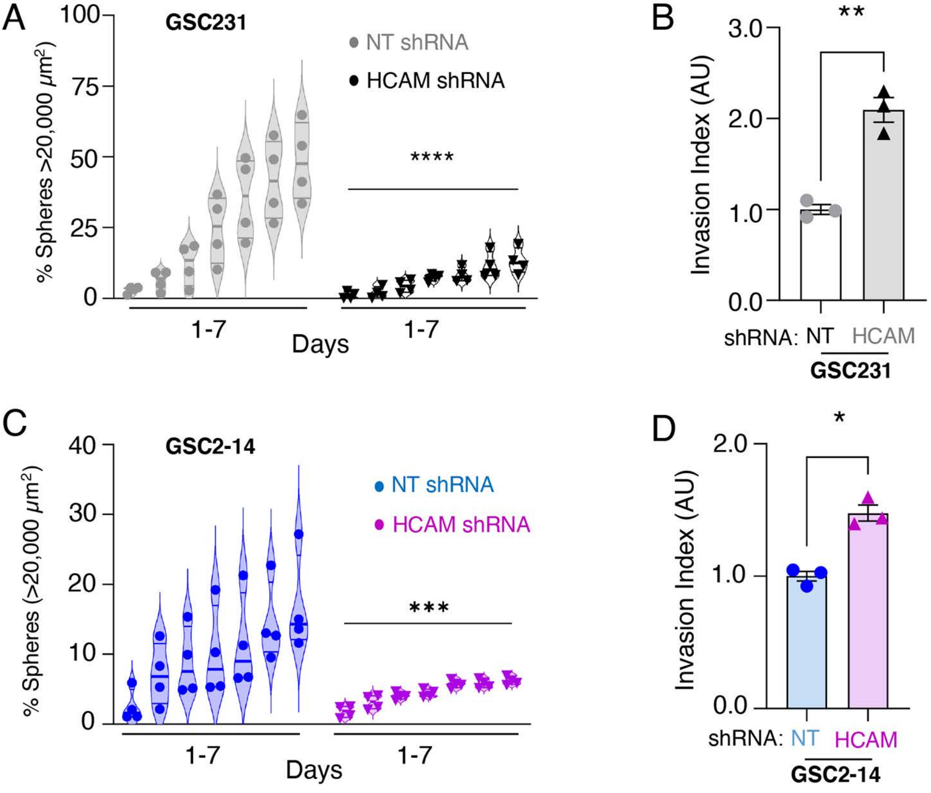

### Supplemental Figure 9

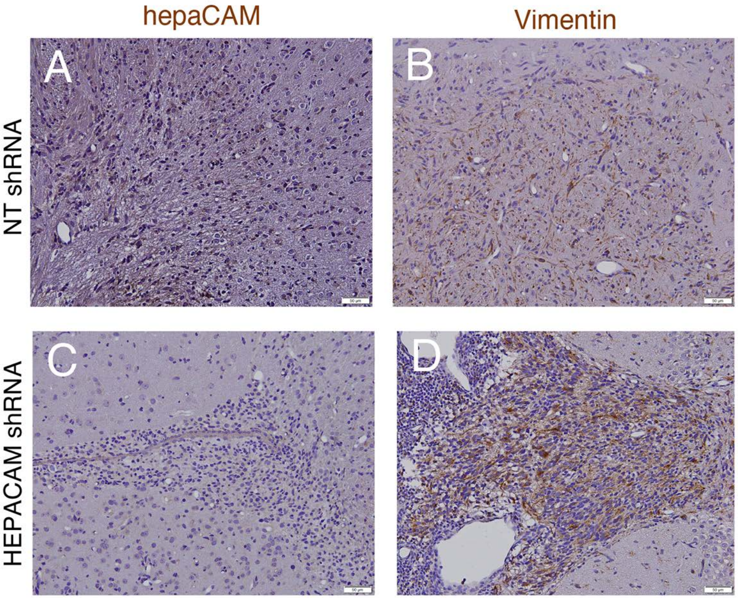
